# Mitochondrial dsRNA from B-ALL cells stimulates mesenchymal stromal cells to become cancer associated fibroblasts

**DOI:** 10.1101/2023.09.26.559490

**Authors:** Richard J. Burt, Aditi Dey, Ayse Akarca, Hermione Allen, Rodothea Amerikanou, Samantha Atkinson, David Auty, Jenny Chatzigerou, Emily Cutler, Jose Afonso Guerra-Assuncao, Kristina Kirschner, Ruchi Kumari, Jiten Manji, Teresa Marafiotti, Adele K. Fielding

## Abstract

Cancer associated fibroblasts (CAF) arising from bone marrow-derived mesenchymal stromal cells (MSC) are prominent in B-precursor acute lymphoblastic leukaemia (B-ALL). We have previously shown that CAF formation is triggered by exposure to reactive oxygen species-inducing chemotherapy and that CAF support chemoresistance by donating mitochondria to the cancer cells, through tunnelling nanotubes. In the present study, we show that exposure of MSC to ALL cell lines, PDX and primary cells or their conditioned media can also trigger CAF formation, in an oncogene-dependent manner. Using bulk RNA sequencing in cell lines, we show that the MSC to CAF transition is accompanied by a robust interferon pathway response and we have validated this finding in primary cells. Using confocal microscopy and flow cytometry, we identify the take up of leukaemia cell-derived mitochondrial dsRNA by MSC as a proximate trigger for the MSC to CAF transition. We show that degradation of dsRNA in ALL cell conditioned media by DMSO ablates the ability of the conditioned media to stimulate MSC to CAF transition. Since we find that only specific primary driver genetic subtypes of B-ALL possess the property to directly generate CAFs, we propose this phenomenon as the first mechanistic insight into the strong relationship between acute lymphoblastic leukaemia genetic subtype and survival outcomes.

**Key points:** X

Exposure of MSC to B-precursor ALL cell lines triggers cancer-associated fibroblast formation in an oncogene-dependent manner
The proximate trigger for CAF formation is ALL-derived mitochondrial double stranded RNA

## Introduction

The primary genetic driver lesion is strongly prognostic for survival in B cell precursor acute lymphoblastic leukaemia (B-ALL)(1) but there is no known mechanism by which genetic driver lesions influence outcome. We have previously demonstrated that an activated mesenchymal stromal cell (MSC) niche develops in response to reactive oxygen-species (ROS)-inducing chemotherapy. The activated MSC that we identified had a cancer associated fibroblast (CAF) phenotype (2) and actively transferred mitochondria along tunneling nanotubes to ‘rescue’ B-ALL cells from chemotherapy, resulting in chemoresistance(3). Dobson and colleagues recently identified a proportion of B-ALL cases in whom chemoresistant sub-clones exist prior to chemotherapy(4), stimulating us to evaluate whether CAF are also present prior to chemotherapy treatment. Here we demonstrate, using a variety of models, that specific genetic subtypes of B-ALL can directly generate CAF from MSC in the absence of chemotherapy. We identify transfer of mitochondrial dsRNA from cancer cells to MSC as a novel mechanism by which MSC can become CAF.

## Methods

### Cells

#### Primary cells

Primary ALL cells and MSC from patients were sourced from participants enrolled on the UKALL14 trial NCT01085617. Normal MSCs were sourced from healthy individuals undergoing bone marrow harvest as donors. All human material was used with informed consent, according to the Declaration of Helsinki.

#### Isolation and Expansion of Primary Human MSC

Mononuclear cells (MNC) were isolated from fresh bone marrow specimens by density gradient centrifugation (Ficoll, Amersham Biosciences, Bucks, UK). Mesenchymal stromal cells (MSC) were isolated and expanded in Mesencult MSC basal medium (STEMCELL^TM^ Technologies) supplemented with Mesencult stimulatory supplements (STEMCELL^TM^ Technologies), 100 units/ml penicillin G (Gibco), 100mg/ml streptomycin (Gibco), 2mM L-glutamine (Gibco) and 1ng/ml basic fibroblast growth factor (R&D Systems). MSC used in experiments were from passage 4 – 5. MSCs were characterized based on the criteria set out by the International Society for Cellular Therapy (ISCT)^(ref)^ using the Human Mesenchymal Stem Cell Functional Identification Kit (R&D Systems) and Human Mesenchymal Stem Cell Verification kit (R&D Systems).

#### Cell Lines

The human MSC cell line HS27a (ATCC); B-precursor ALL cell lines REH, SD1, SEM, and TOM1; and the murine MSC cell line MS5, and the cervical cancer cell line HeLa (all from DSMZ) were grown in RPMI 1640 (MS5, αMEM) (HeLa, DMEM) with 5% to 20% fetal bovine serum and penicillin/streptomycin/glutamine.

#### Exposure of MSC to ALLs or media

For co-culture experiments, MSC cell lines were plated on Day 0, and ALL cells at a ratio of 1:4 were added on Day 1. The cells were flow-sorted after 3-5 days and subjected to imaging or nucleic acid extraction. For transwell experiments, the ALL cells were added onto a transwell insert (0.1-0.4μm) (Greiner Bio-one) at Day 1. For contact-independent experiments, MSCs were incubated with the B-ALL cell conditioned medium.

#### Mitochondrial depletion

SD1 cells were cultured in media containing 0.1µg/ml Ethidium Bromide supplemented with uridine 50 µg/ml and 1mM sodium pyruvate for 4 weeks. Mitochondrial depletion was confirmed by PCR and imaging.

#### DMSO degradation of dsRNA from conditioned media

Conditioned medium was treated with 1%DMSO overnight, followed by quantification of dsRNA by ELISA (Exalpha/ Nordic Mubio) to confirm depletion

### Microscopy

#### Immunocytochemistry

Immunocytochemistry was performed according to a modified R&D Systems protocol. Cells were fixed with 4% paraformaldehyde, washed and then blocked for 2 hours with a buffer containing 1% bovine serum albumin (BSA), 10% normal donkey serum (Abcam) and 0.3% Triton X-100 (Sigma Aldrich). After blocking, the nuclear stain DAPI (Santa Cruz Biotechnology) and F-actin stain Phalloidin-Atto 633 (Sigma Aldrich) stain were added for 10 minutes. Imaging was done with a Zeiss axio-observer Z1 fluorescent microscope.

#### Immmunofluorescence microscopy for dsRNA detection

Adherent cells were stained directly in 24-well, glass bottom plates, while B-ALL suspension cells were adhered to fibronectin-coated, 24-well glass bottom plates overnight before staining. Cells were stained with MitoTracker™ Deep Red FM (Invitrogen™) according to manufacturer’s instructions. The cells were then washed with PBS, fixed with 4% paraformaldehyde and permeabilised with 0.25% Triton^TM^ X-100 (Sigma-Aldrich) in PBS.

After washing with 0.05% Tween-20 in PBS, 3% BSA (BSA-heat shock fraction - Sigma Aldrich) was added as a blocking agent for 1 hour. After blocking, the anti-dsRNA antibody J2 (SCICONS) was added at 1:200 dilution in 3% BSA followed by the DyLight® 488 Anti-Mouse IgG (H+L) secondary antibody (2BScientific) (1:300 dilution) for one hour each. Finally, the nuclear stain DAPI (Santa Cruz Biotechnology) was added for 10 minutes. Imaging was done using the LSM880 confocal microscope.

#### Immunohistochemistry of femur sections

After fixation in 10% neutral buffered formalin, samples were decalcified in 10% formic acid for 9-10 hours then processed and embedded and sectioned. Samples were stained with anti-human CD19 and anti-mouse nestin (Abcam) primary antibodies.

### Flow Cytometry

#### Flow Cytometric detection of dsRNA

Cells were washed and resuspended in PBS. They were then incubated with fixable viability dye Zombie NIR^TM^ dye (BioLegend) for 15 minutes at 4 degrees. The cells were then incubated with anti-human CD19 (BD Biosciences) for 30 minutes at 4 degrees before fixation and permeabilisation with 4% PFA and 0.1% Triton^TM^ X-100 (Sigma-Aldrich) respectively. The cells were blocked in 1% BSA (BSA-heat shock fraction - Sigma Aldrich) for an hour at room temperature. After blocking, anti-dsRNA antibody J2 (1ug/ml) (SCICONS)was added at 1:100 dilution in 1% BSA followed by DyLight® 488 Anti-Mouse IgG (H+L) secondary antibody (2BScientific) (1:300 dilution) for one hour each. The cells were washed and then run on BD LSRFortessa^TM^ X-20. Data were analysed with FLOWJO.

#### Fluorescent Activated Cell Sorting of HS27a MSC

Following the co-culture of ALL cells and MSCs for five days, the MSCs and ALL cells were removed from the plate by trypsinisation and sorted on expression of CD90 using anti-CD90 FITC or anti-CD90 APC (BD Biosciences) and lack of expression of CD19 using anti CD19 APC or CD19 BV605 (BD Biosciences) using BD FACSAria^TM^ III cell sorter.

#### Mitotracker assay

MSC were stained with MitoTracker^TM^ Red (ThermoFisher M22426) according to the manufacturer’s instructions at 37°C for 30 mins. The cells were washed twice, then left for three hours to eliminate unbound probe prior to a final wash. The stained MSC were co-cultured with different ALL cell lines or primary ALL cells for 24 – 72 hours. The ALL cells were then collected and stained with antiCD19 BV605 (BD Biosciences) followed by analysis for the presence of mitochondria in the CD19-positive ALL cell population.

#### Quantification of reactive oxygen species

B-ALL cell lines were collected and stained with CellROX^Ⓡ^Green (ThermoFisher C10444) according to manufacturer’s instruction and their ROS levels measured by flow.

### Quantification of secreted proteins

#### Cytometric bead array

A cytometric bead array (BD Biosciences) was used according to the manufacturer’s instructions with the IL6 Flex Set (558276; BD Biosciences), human IL8 Flex Set (558277; BD Biosciences), or human MCP-1/CCL2 Flex Set (558287; BD Biosciences). Three hundred events per analyte from the live gate were collected on a BD FACSAria instrument (Becton Dickinson). Data were analyzed with FCAP Array Software, version 3.0.

#### Enzyme-linked immunosorbent assay

The supernatant was collected from the MSCs at the end of the experiment (48 to 72hr). VeriKine-HS Human Interferon Beta TCM ELISA Kit 96T (PBL Assay Science) was used to quantify IFNß in the supernatant.

A double-stranded RNA ELISA kit (Exalpha/Nordic Mubio) was used to quantify dsRNA from 1ug of total RNA, according to manufacturer’s instructions.

### Molecular Biology

#### RNA extraction

RNA was extracted from cells with TRIzol (15596026; Ambion, Life Technologies) and separated from DNA by using chloroform (Sigma-Aldrich). Isopropanol (Sigma-Aldrich) was added, and the samples were frozen overnight at −80°C. After it was thawed and washed with 70% ethanol, the pellet of RNA was resuspended in RNase-free water, and the concentration was measured on a NanoDrop spectrophotometer. RNA was also extracted using the RNeasy Micro Kit (Qiagen) or RNeasy Mini Kit (Qiagen) according to the manufacturer’s protocol.

#### RT2 profiler PCR assays

cDNA was synthesized with the RT2 First Strand kit (330401; Qiagen), according to the manufacturer’s instructions. The cDNA was then used for an RT2 Profiler PCR array, according to the manufacturer’s protocol with a predefined and pre prepared selection of primers for appropriate CAF-defining targets or RNA sensing gene targets listed in supplemental Tables 1 and 2. Each sample was run in triplicate for each gene and quantified relative to the glyceraldehyde-3-phosphate dehydrogenase housekeeping control. For the RNA sensing panel, the experiments were done in triplicate but the samples were run once. *Mitochondrial DNA detection*

DNA was extracted from cells using QIAamp® DNA Blood Mini Kit (Qiagen). The DNA was amplified for detection of mitochondrial and nuclear DNA from both human and mouse using the primers below. Annealing temperature used was 60 degrees for 15 to 25 cycles. The PCR products were run in 2% agarose (SIGMA) gels and visualised under UV light.

#### RT-PCR for mitochondrial dsRNA immunoprecipitated from B-ALL conditioned media

dsRNA was immunoprecipitated using Protein G Dynabeads (Thermofisher). The protocol was modified from the manufacturer’s method. Briefly, 5ug of J2 mAb was coated onto protein G magnetic dynabeads and then the SD1/SEM conditioned media was incubated with the J2 mAb coated dynabeads for 2hrs at 4 degrees. The beads were then centrifuged and washed before RNA was extracted using TriZol method. The RNA was then subjected to reverse transcriptase PCR to obtain cDNA. Once cDNA was made, it was used to perform an end point qualitative PCR using the following primers. Total RNA extracted from SD1 cells were used as a positive control.

#### RNA sequencing

HS27a cells were plated and incubated with SD1 or SEM conditioned media or RPMI control in triplicate and RNA was extracted from the MSCs 48-72hr later. RNA from experimental triplicates was processed and sequenced by the GOS UCL Genomics Facility (London, UK).

Libraries were prepared using the KAPA mRNA Hyper Prep kit. Samples were sequenced on the Illumina NextSeq 2000. FastQ files were assessed with FastQC (Babraham Institute Bioinformatics Group). Adapter sequences, primers, poly-A tails were removed with Cutadapt. Raw data was mapped with the STAR aligner to the human genome GRCh38 with annotation from Ensembl. Gene expression analysis was carried out in R (version 4.2.2).

Differentially expressed genes between control and exposed HS27a were determined using DeSEQ2(14). Significantly upregulated genes were determined using a p-adjusted value < 0.05. Gene set enrichment analysis (GSEA) was performed with fgsea using the Reactome pathway sets(15) as reference. The significance of enriched pathways was determined using a p-adjusted value of <0.05.

### Mouse experiments

All animal experiments were performed according to UK Home Office approved protocols and institutional guidelines. Disseminated SEM/SD1 xenografts were established in 8-10-week-old NSG (Non obese diabetic severe combined immunodeficiency gamma) male mice (Charles River, Margate, UK) by tail vein injection of 2 x 10^6^ SEM or SD1 cells expressing luciferase and blue fluorescent protein. A mouse without leukaemia cells was used as a baseline control for imaging. Engraftment was confirmed after 7 days and disease was monitored by bioluminescent imaging; mice were shaved, injected i.p. with 200 μl of D-luciferin (Caliper Life Science, Cheshire, UK) and imaged under isofluorane anaesthesia in an IVIS 100 Lumina (Caliper Life Sciences, Chesire, UK). The data were analyzed using Living Image 3.2 software. Mice were sacrificed at the humane end point, and a femur was fixed in 10% neutral buffered formalin for immunohistochemistry.

### Statistical analysis

The data generated were analysed on GraphPad Prism 6 software or other software where indicated. For statistical comparison Chi-squared or unpaired Student t tests were used, as indicated.

## Results

First, we sought to determine if B-ALL cells could directly activate MSC to become CAFs. We co-cultured five distinct B-ALL cell lines as well as control, healthy donor B-cells together with HS27a, a human MSC cell line. We used our previously published criteria(3) to assess CAF formation, namely, cytoskeletal changes (broadened and flattened morphology with prominent actin stress fibres), upregulation of relevant genes in our targeted 18-gene MSC activation/CAF qPCR panel and IL6, IL8 and CC2 secretion. Three of five B-ALL cell lines; *697 (TCF3::PBX), TOM-1, SD1 (*both *BCR::ABL1)* ) induced characteristic CAF changes in the HS27a cells (Figure 1a). REH (*ETV6::RUNX1*) cells did not induce CAF-like morphological changes in the HS27a cells but did upregulate a selection of CAF-relevant genes. SEM (*KMT2A::AFF4*) cells and healthy B-cells induced neither morphological changes nor upregulation of CAF-relevant genes in the HS27a cells. A cytokine bead array (Figure 1b) showed a significant increase in IL6, IL8 and CCL2 proteins in the media of SD1 cells co-cultured with HS27a as compared to mono-culture or SEM cells in co-culture, in keeping with upregulation of relevant genes in the qPCR data. We confirmed that CAF-induction also occurred after co-culture of primary patient ALL cells with normal, healthy donor MSC obtained from two healthy bone marrow donors. Three of four primary B-ALL samples [14-1-285 (*BCR::ABL1*), 14-1–456 (B-other, no known primary driver lesion), 14-1-475 (*BCR::ABL1*), 14-1-479 (*BCR::ABL1*)] tested induced activation of two different healthy donor MSC, as characterised by cytoskeletal changes (figure 1c) as well as IL6, IL8 and CCL2 secretion (Figure 1d). Given our own and others’ findings that CAF can be induced by exposure to ROS, we quantified the intrinsic ROS level of the cell lines by CellROX© assay. Figure 1e shows the relationship between B-ALL cell intrinsic ROS level of the cell as measured by CellROX© and CAF-generating capacity. We further confirmed that the MSC activation without chemotherapy occurred in a murine xenograft model; we selected SD1 cells and SEM cells for the model due to their divergent abilities to induce CAF *in vitro*. Luciferase-expressing SD1 and SEM cells were injected via the tail vein (N=4 mice per group) and, 3 days after engraftment was confirmed at 14 days, the mice were sacrificed and femora dissected. Femoral sections stained with antibodies against CD19 and nestin showed a striking preponderance of nestin-expressing cells in the SD1 model than the SEM model, consistent with *ab-initio* CAF formation; a representative example is shown in figure 1f.

**Figure 1.**
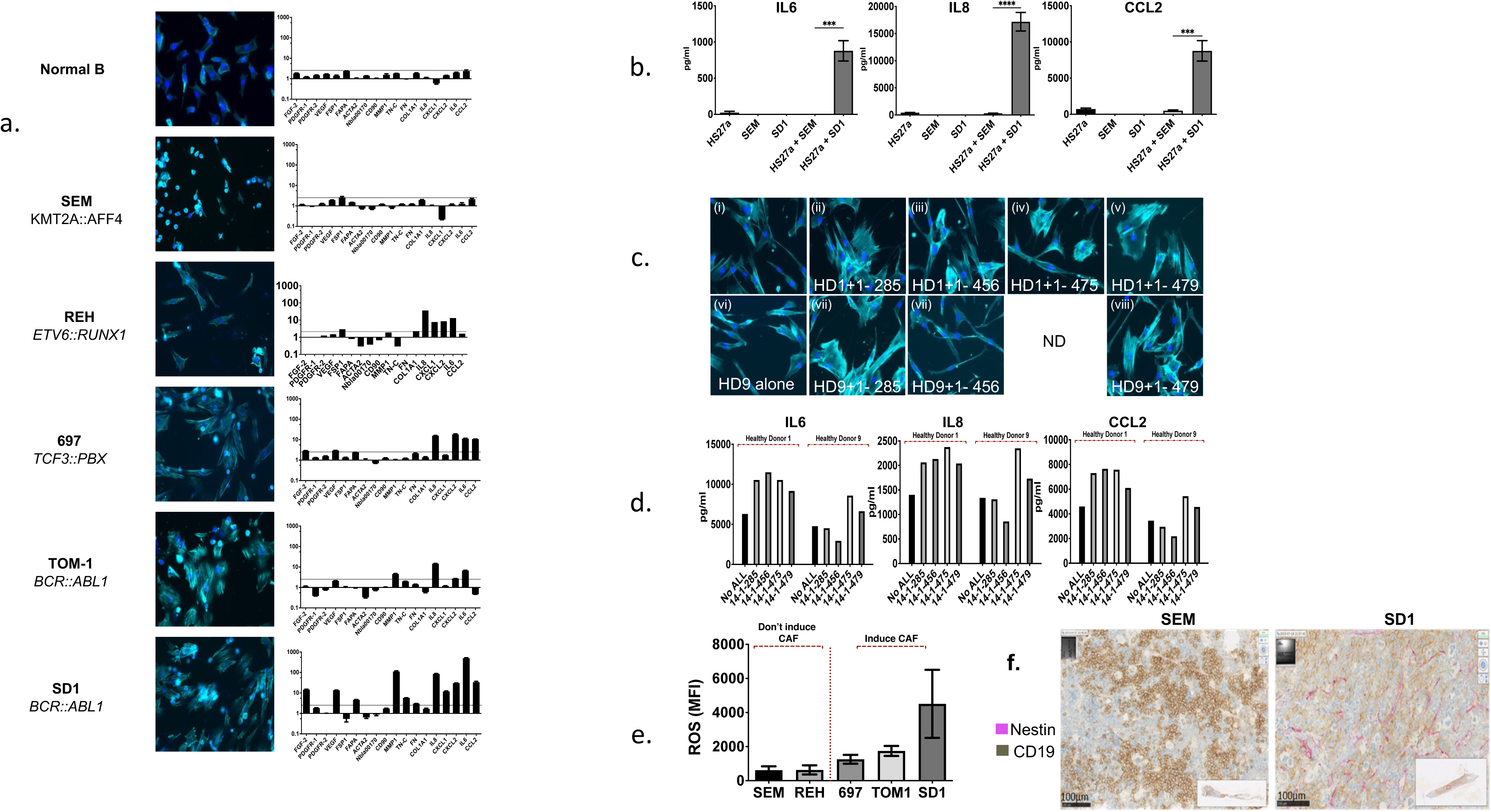
B precursor acute lymphoblastic leukaemia cells directly stimulate mesenchymal stromal cells to become cancer-associated fibroblasts. **a.** Photomicrographs (20x magnification) showing showing phalloidin/DAPI staining of HS27a MSC after co-culture with a series of ALL cell lines of different genetic subtypes, as indicated, alongside a gene expression panel of showing fold upregulation (Y axis) of the 18 gene CAF panel compared to mean baseline of HS27a MSC. **b.** Cytokine bead assays for IL6, IL8 and CCL2 (pg/ml, Y axis) following co-culture of HS27a MSC with SEM or SD1 ALL cells and controls of each cell alone, as indicated on the X axis. Mean and standard error of mean from 3 independent experiments are shown. P values for comparisons between HS27a + SEM and HS27a +SD1 by unpaired t-test are 0.0004 (IL6), <0.0001 (IL8) and 0.0005 (CCL2) **c.** Photomicrographs showing phalloidin/DAPI (40x magnification) staining of two normal healthy donor (HD1, HD9) MSCs after co-culture with 4 different primary patient ALL samples, 1-285, 1-456, 1-475 and 1-479. **d.** IL6, IL8 and CCL2 (pg/ml, Y axis) secreted by HD1 and HD9 MSC after co-culture with the 4 individual primary patient ALL samples **e.** Mean fluorescent intensity (Y axis) of reactive oxygen species after CellROX^Ⓡ^Green staining of the panel of ALL cells lines used in a. Mean and standard error of mean from 3 independent experiments are shown. **f.** Representative sections of femur from NSG mice with established leukaemia derived from SD1 or SEM cells stained by CD19 (brown) or Nestin (pink)

We sought to elucidate the mechanism behind the differential CAF induction by B-ALL cells by different genetic subtypes. First, we showed that cell-cell contact was not required; SD1 but not SEM cells still induced CAFs from HS27a when they were co-cultured but separated in a transwell (data not shown). SD1 but not SEM conditioned media similarly induced CAFs, as shown by the imaging and RQ-PCR (figure 2a) as well as cytokine bead assays ( figure 2b). A Proteome Profiler Human XL cytokine array assay for cytokines/chemokines in the conditioned media of SD1 and SEM cells did not reveal any likely causative cytokine candidates. In view of the relationship shown in figure 1 between intracellular ROS levels CAF-induction, we asked the question if mitochondria were transmitted from B-ALL cells to MSC, as mitochondria are an important source of ROS within the cell. A flow cytometric mitotracker dye assay (figure 2c) showed an approximate three-fold higher transfer of mitochondria from SD1 than SEM cells. To rule out passive dye transfer as responsible for the flow cytometric findings, we co-cultured the murine MSC cell line MS5 with human leukaemia cells, then sought the presence of human mitochondrial DNA in the flow-sorted murine MS5 cells. These data are shown in fig 2d and confirm human mitochondrial - but not nuclear - DNA in the mouse stroma. To confirm that SD1 mitochondria were associated with CAF induction in HS27a we repeated the imaging and cytokine release CAF assays after depletion of mitochondrial nucleic acid from the SD1 cells by a lengthy co-culture with low dose ethidium bromide(5). Fig 2e shows that SD1 cells depleted of mitochondrial nucleic acid were less able to induce CAF-like morphological changes and generated significantly lower levels of IL8 and CCL2, although not IL6, secretion by HS27a MSC. However, on exposing HS27a to SD1-conditioned medium having filtered out whole mitochondria (figure 2f) using a 0.2µm filter, we did not abrogate IL6, IL8 or CCL2 production at all, suggesting that CAF induction results from transfer of mitochondrial fragments or mitochondrial nucleic acid.

**Figure 2.**
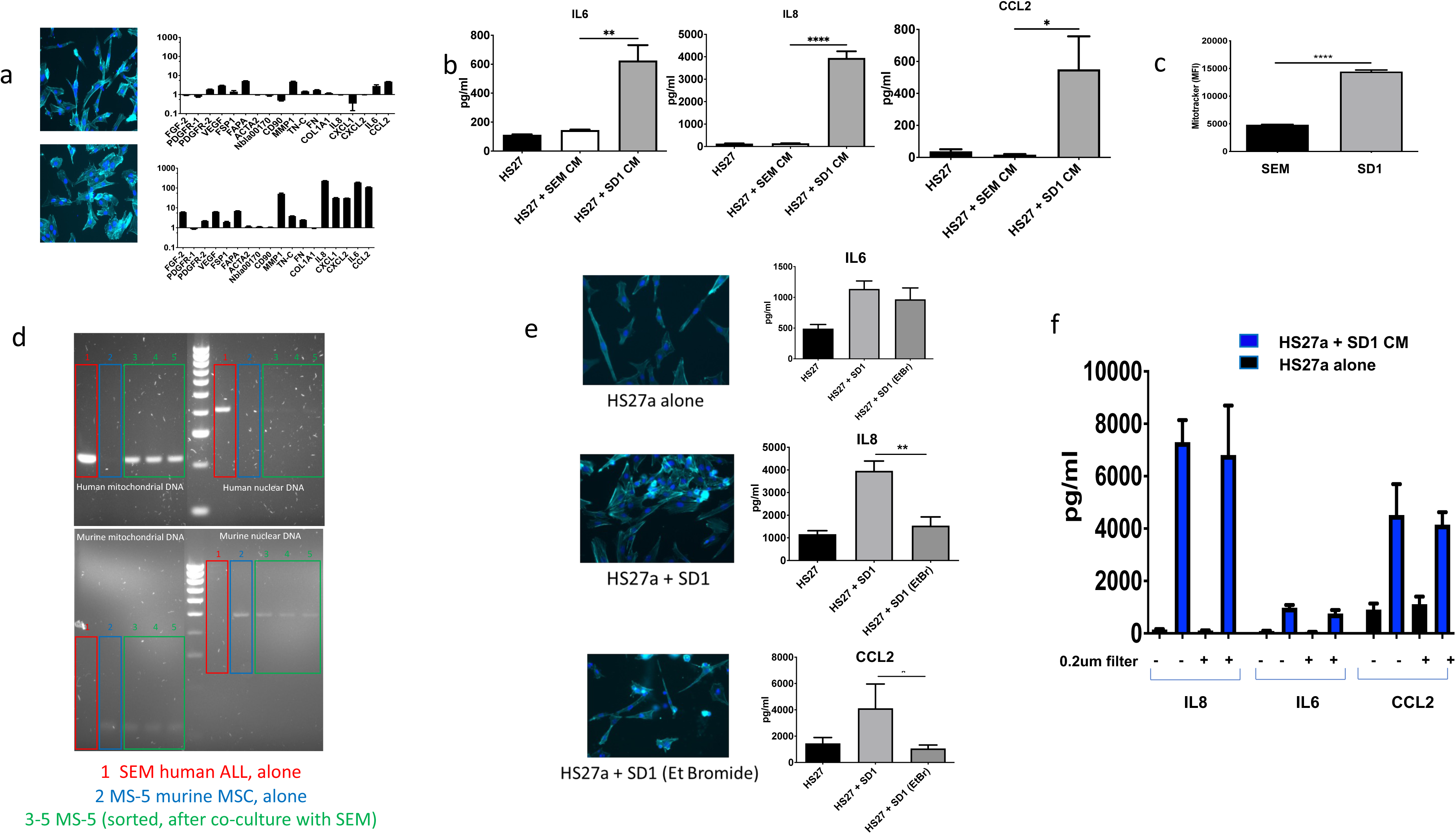
The B precursor ALL-mediated mesenchymal stromal cell to cancer associated fibroblast transition is contact-independent and is mediated by transfer of mitochondrial elements. **a.** Photomicrographs (20x magnification) showing showing phalloidin/DAPI staining of HS27a MSC after co-culture with SEM (top panel) or SD1 (bottom panel) conditioned media alongside the 18 gene panel showing fold upregulation (Y axis) compared to mean baseline of HS27a MSC **b.** Cytokine bead assays for IL6, IL8 and CCL2 (pg/ml, Y axis) following exposure of HS27a MSC to RPMI alone or SEM or SD1 ALL conditioned media, as indicated on the X axis. Mean and standard error of mean from 3 independent experiments are shown. *.01 < P ≤ .05; **.001 < P ≤ .01; ****P ≤ .0001 **c.** Mitochondrial transfer to HS27a MSC from SEM or SD1 cells (X-axis) by mitotracker assay, (mean fluorescence intensity, Y axis), from HS27a to SEM cells **d.** Agarose gel images showing PCR products from human nuclear and mitochondrial DNA and murine nuclear and mitochondrial DNA - as indicated in each quadrant, - after co-culture of SEM cells with MS5 murine stromal cells. Lane 1 Red boxes SEM alone, Lane 2 Blue boxes MS5 alone, Lanes 3-5 Green boxes MS5 cells flow sorted coculture after with MS-5. **e.** Photomicrographs (20x magnification) showing showing phalloidin/DAPI staining of HS27a MSC alone and cytokine bead assays for IL6, IL8 and CCL2 (pg/ml, Y axis) production by HS27a cells; alone, after co-culture SD1 cells and after coculture with SD1 cells depleted of mitochondrial nucleic acid by low dose ethidium bromide. For the cytokine production, mean and standard error of mean from 3 independent experiments are shown **f.** Cytokine bead assays for IL6, IL8 and CCL2 (pg/ml, Y axis) following exposure of HS27a cells to SD1-conditioned medium (blue bars) or RPMI alone control (black bars) with (+) or without (−) 0.2µm filtration.

In order to gain a more global understanding of the phenomenon, we performed bulk RNA sequencing on HS27a exposed to CAF-inducing and non-inducing conditions. We used conditioned media from SD1 and SEM cells to rule out any contamination by B-ALL cells. RPMI alone served as a baseline. Figure 3a shows the volcano plots of HS27a cells grown in either SD1 or SEM conditioned media, each compared to RPMI control. Of specific interest, the most highly upregulated genes in HS27a after exposure to SD1 but not SEM conditioned media, were mostly genes encoding cytokines and chemokines (*CCL5, CXCL10, CXCL11, CCL2, CXCL1*), RNA sensing genes (*OAS1, OAS2, OAS3, MX1*) or interferon pathway related genes (*IFI44L, IFIH1, IFIT3*). Gene enrichment pathway analysis revealed that the most enriched pathways in the SD1 conditioned medium condition were related to interferon signalling as seen in Figure 3b. Other pathways which were enriched included cytosolic nucleic acid sensing, cytokine/chemokine signalling, and TNF⍺ signalling. Figure 3c shows the most highly-upregulated interferon pathway genes in SD1 conditioned media compared to RPMI. Strikingly, many of these genes are involved in RNA sensing or inhibition (*MX1-2, OAS1-3, IFIH1/MDA-5, DDX58/RIG-I*). We measured the levels of interferon α,β and γ by ELISA in the media of HS27a cells exposed to SD1 or SEM conditioned media or RPMI. We did not see any secreted IFN α or γ in any of the conditions, although we detected very low amounts (maximum level of 68pg/ml) of IFNβ (data not shown) produced by SD1- conditioned media exposed-HS27a at the 72 hour time point, ruling out a classic antiviral response. We used RT-PCR to validate the findings by in the 2 different normal healthy donor MSC samples after exposure to the same primary ALL samples (figure 3d) that were used in figure 1.

**Figure 3.**
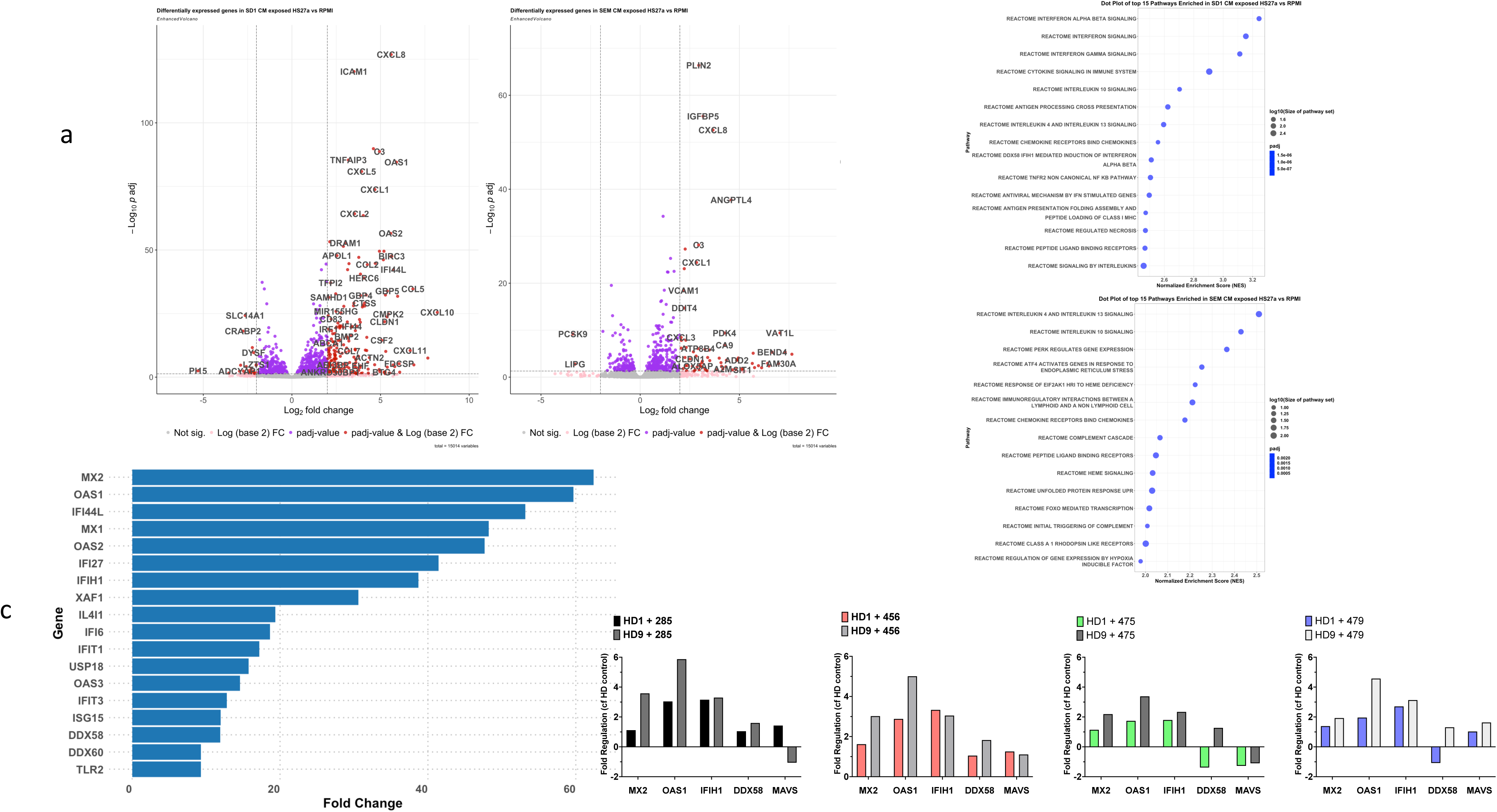
The B precursor ALL-mediated MSC to CAF transition is accompanied by a robust interferon pathway response. **a.** Two volcano plots showing differentially-regulated genes from HS27a cells cultured in SD1 conditioned medium compared to RPMI andHS27a cells cultured in SEM conditioned medium compared to RPMI, as indicated. **b.** Dot plots of the top 15 pathways enriched in HS27a cells cultured in SD1 conditioned medium compared to RPMI and HS27a cells cultured in SEM conditioned medium compared to RPMI, as indicated. Dot size is proportional to log_10_(size of pathway set) and colour is proportional to adjusted p value. All p values were highly significant, as indicated. **c.** Bar chart showing fold upregulation (X axis) of interferon pathway genes indicated on the Y axis. **d.** Validation of RNAseq interferon response finding in two primary, healthy donor (HD1 and HD9) MSCs co-cultured with four different primary ALL cell specimens (X axes). Y axis indicates fold change in gene expression.

We were prompted by a recent finding that mitochondrial dsRNA can generate an antiviral signalling response in humans(6) to explore mitochondrial dsRNA as the cause for CAF formation. We sought the presence of mitochondrial dsRNA in the conditioned medium of ALL cells as an explanation for our findings. We co-stained SD1, SEM and HS27a cells alone as well as HS27a exposed to SD1 and SEM conditioned medium with the J2 dsRNA-specific antibody, deep red mitotracker and DAPI then carried out confocal microscopy. HeLa cells served as a positive control. Figure 4a shows very prominent dsRNA staining in SD1 cells as well as in HS27a cells exposed to SD1 conditioned media but almost complete absence of dsRNA in SEM cells and HS27a cultured in SEM conditioned media or RPMI. To confirm these findings with a different technique, we measured MFI by flow cytometry after J2 staining, using the same experimental conditions. As shown in figure 4b, there was an approximate 1.5-fold increase in the MFI of dsRNA expression by flow cytometry in HS27a exposed to SD1 conditioned media compared to SEM conditioned media or RPMI. Next, to confirm that the dsRNA in the SD1 was indeed of mitochondrial origin, we immunoprecipitated dsRNA from the SD1 and SEM conditioned media using the J2 antibody. Total cellular RNA from SD1 served as a positive control. After synthesis of cDNA, we performed PCR analysis of four mitochondrial genes mtCOX1, mtND5 and mtND6 and mtCYB, as well as the nuclear gene GAPDH. Figure 4C shows amplification of mitochondrial genes after J2-immunoprecipitation of SD1 but not SEM conditioned medium. We confirmed the presence of dsRNA (% cells stained with J2 antibody by flow cytometry) in whole bone marrow and then CD19-sorted cells taken from three *BCR::ABL1* (1,4,6) and three *KMT2A::AFF4* (2,3,5) patient derived xenograft (PDX) samples grown in NSG mice (figure 4d). NSG mouse marrow alone was the negative control. We found dsRNA in a percentage of cells in all the specimens but as the experiment was only done once, we cannot make inferences about the comparative levels.

**Figure 4.**
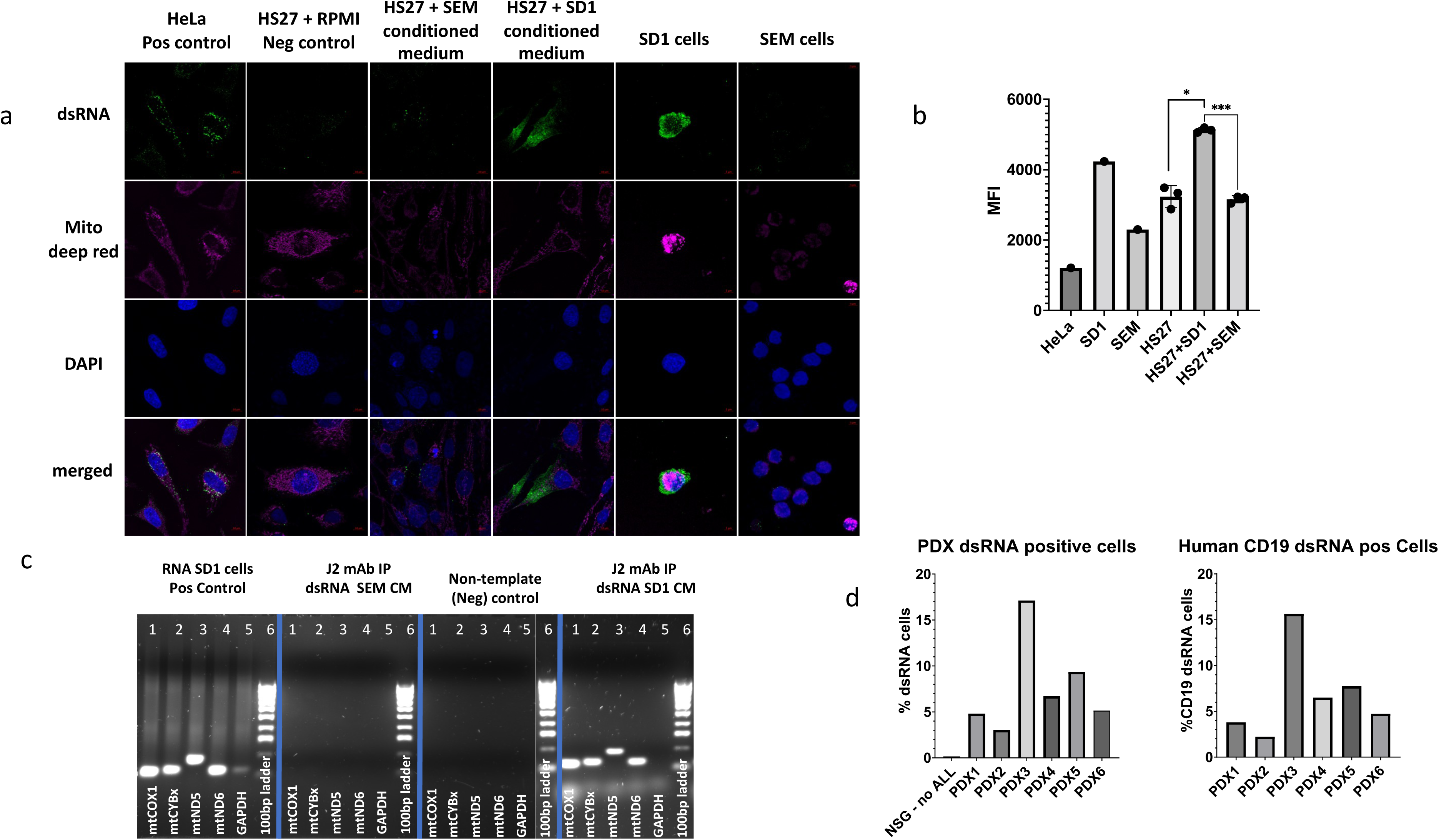
CAF-inducing B-precursor ALL cells contain mitochondrial double-stranded RNA which is released and taken up by MSC. **a.** Photomicrographs of confocal images of cells stained with anti-dsRNA monoclonal antibody J2, deep red mitotracker and DAPI as well as merged images, as indicated on the left. Cell types and experimental conditions are indicated above the image. **b.** Mean fluorescence intensity (MFI, Y axis) after intracellular J2 staining and flow cytometry of the cells indicated on the X axis. HS27a+SD1 or SEM indicates HS27a cells exposed to the SD1 or SEM conditioned medium **c.** Agarose gel images showing the products of RT-PCR for four mitochondrial genes, and GAPDH with a 100 base pair ladder (as indicated below the bands) from the experimental conditions indicated at the top of the individual gel images: positive control (SD1 cells), J2-immunoprecipitated RNA from SD1 and SEM cells and negative non-template control. **d.** Bar chart showing percentage of ALL PDX cells (Y axis) positive for dsRNA after intracellular J2 staining and flow cytometry. PDX 1,4 and 6 are derived from *BCR::ABL1*+ ALL and PDX 2,3 and 5 from *KMT2A::AFF4*+ ALL

In order to demonstrate the key role of mtdsRNA in CAF formation, we used DMSO to degrade the dsRNA(7) from the conditioned medium of SD1 cells. As shown in figure 5a, An ELISA for dsRNA carried out on SD1 conditioned medium quantified a mean baseline dsRNA concentration of around 1200 pg/ml which was reduced to the limit of detection by 1% DMSO. DMSO-treated SD1 conditioned medium was compared to regular conditioned medium (baseline, set at 1) for the ability to upregulate CAF (figure 5b) and RNA sensing genes (figure 5c). Figure 5 b and c show down regulation of both those gene set after DMSO-treatment compared to the baseline condition. Figure 5d shows CAF-associated cytokine secretion after DMSO-treated SD1 conditioned medium; IL6, IL8 and CCL2 secretion were all significantly reduced by degradation of dsRNA from the conditioned medium.

**Figure 5.**
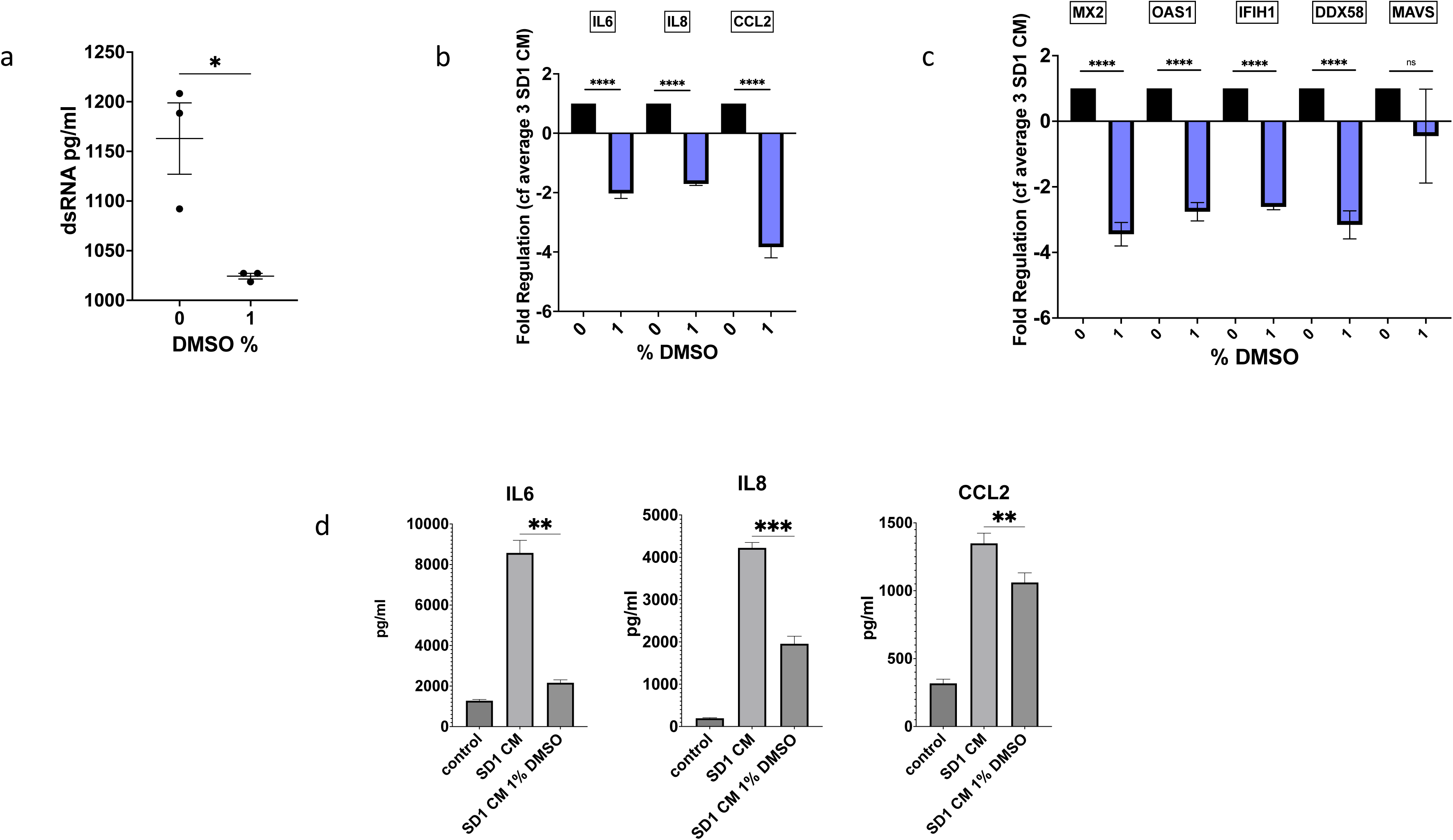
Degradation of mitochondrial dsRNA prevents the interferon response and the MSC to CAF transition. **a.** ELISA quantification of dsRNA (pg/ml, Y axis) from SD1 conditioned medium with and without 1% DMSO treatment. The lines denote the mean and error bars SEM of 3 independent experiments. The difference with and without DMSO is statistically significant, p=0.0183. **b.** Fold regulation of CAF gene expression (Y axis) in HS27a cells exposed to SD1 cell conditioned medium (set at 1) denoted by 0 on the X axis compared to HS27a cells exposed to DMSO-treated SD1 cell conditioned medium denoted by 1 on the X axis. Bars show mean and SEM of 3 independent experiments. All p values <0.0001. **c.** Fold regulation of expression of four interferon pathway genes expression (Y axis) in HS27a cells exposed to SD1 cell conditioned medium (set at 1) denoted by 0 on the X axis compared to HS27a cells exposed to DMSO-treated SD1 cell conditioned medium denoted by 1 on the X axis. Bars show mean and SEM of 3 independent experiments. All p values are p<0.0001. **d.** Cytokine bead assays for IL6, IL8 and CCL2 (pg/ml, Y axis) following exposure of HS27a MSC to RPMI alone, SD1 conditioned medium, or SD1 conditioned medium treated with 1% DMSO, as indicated on the X axis. Mean and standard error of mean from 3 independent experiments are shown. The p values are p**=00021 (IL6); p***=00003 (IL8); p**=0.0039 (CCL2).

## Discussion

We have identified a novel biological mechanism by which exposure to leukaemia-cell derived mitochondrial dsRNA can generate CAF from MSC in the absence of chemotherapy.

We revealed a differential capacity to generate CAF which was associated with the intrinsic ROS level of the cancer cell. It is already known that oxidative stress, which results from an imbalance between the production of ROS and the ability of cells to scavenge them can trigger the opening of the mitochondrial permeability transition pore which is known to result in extracellular delivery of mtDNA which is normally embedded in the mitochondrial matrix (8, 9). For our subsequent experiments, we focused on *BCR::ABL1+* and *KMT2A::AFF4*+ models of ALL because they were the two genetic subtypes which showed the greatest disparity in both their intrinsic ROS levels and ability to stimulate CAF formation.

Mitochondrial DNA is bidirectionally transcribed, generating overlapping transcripts, which can form long, double-stranded RNA structures whose production is usually restricted by the degradosome components, mitochondrial RNA helicase SUV3 and polynucleotide phosphorylase PNPase (10). However, the recent identification of mitochondrial dsRNA as a generator of an antiviral signalling response in humans(6) coupled with our RNAseq findings showing a strong and very significant interferon pathway upregulation was the proximate trigger for us to focus on mitochondrial dsRNA as a potential trigger for CAF formation. Thereafter, we readily demonstrated dsRNA inside the *BCR::ABL1*+ ALL cells and take-up by stromal cells after exposure to conditioned medium from *BCR::ABL1*+ ALL cells. We also found mtdsRNA in PDX samples grown in NSG mice, although we had insufficient samples to determine a clear relationship with the genetic subtype of ALL. The causal relationship between mtdsRNA and CAF-formation was shown by the abrogation of CAF gene expression and cytokine secretion upon degrading dsRNA in the conditioned media. Our data provide the first mechanistic insight into how different genetic subtypes of ALL differently influence the stromal microenvironment. They pave the way for further studies to determine the extent to which this mechanism contributes to the strong influence of genetic subtype on survival outcomes.

Data from the oncolytic virus field shed light on how transformed cells may ‘tolerate’ the presence of dsRNA whereas the interferon signalling pathway is strongly activated by stromal cells under the same circumstance. It is widely accepted that the particular susceptibility of cancer cells to oncolytic viruses is at least in part mediated by defects in their interferon response in cancer cells as summarised by(11). Previous work from our group used a stepwise model of transformation, in which oncogenic hits were additively expressed in human MSC (12). Untransformed MSC were very resistant to vaccine strain measles virus-mediated oncolysis due to a robust interferon response. However, susceptibility to the virus progressively increased with the level of transformation, due to an increasing delay and very significant reduction in the magnitude of the interferon response. A contact-dependent transcytosis of cytoplasm from cancer cells into fibroblasts which leads to STING and IFR3-mediated expression of interferon-β, subsequently driving interferon-stimulated transcriptional programs in stromal fibroblasts has also been demonstrated (13).

In conclusion, our data reveal a previously unidentified mechanism by which contact-independent take-up by MSC of dsDNA of mitochondrial origin from leukaemia cells drives the MSC to a cancer-associated fibroblast phenotype. Our previous work shows that CAF can protect leukaemia cells from ROS-inducing chemotherapy by donation of mitochondria, demonstrating the clinical relevance of our findings. Studies in other cancer types are warranted.

## Acknowledgements

RB was supported by CRUK Clinician Scientist Fellowship RCCFEL/100017 AD was supported by MRC grant MR/W000148/1 to AKF and KK The UKALL14 trial was funded by CRUK grant CRUK/09/006 to AKF Primary specimens were obtained from UKALL14 biobank which was funded by grant CRUK grant C27995/A21019 to AKF and Anthony V. Moorman

## Author contributions

RB and AD carried out experiments and contributed to manuscript writing. AA, HA RA, SA, JC, EC, RK, KK and TM carried out specific experiments. DA and JG-A provided significant assistance with coding and bioinformatic analyses. JM provided significant assistance with confocal microscopy. AKF conceived, funded and supervised the project and wrote the manuscript. All authors read and commented on the manuscript.

## Disclosures

The authors have no disclosures.

## Notes

### Competing Interest Statement

The authors have declared no competing interest.

